# Acetaminophen inhibits diacylglycerol lipase synthesis of 2-arachidonoyl glycerol: implications for nociception

**DOI:** 10.1101/2025.03.05.640851

**Authors:** Michaela Dvorakova, Taryn Bosquez-Berger, Jenna Billingsley, Natalia Murataeva, Taylor Woodward, Emma Leishman, Anaëlle Zimmowitch, Anne Gibson, Jim Wager-Miller, Ruyi Cai, Shangxuan Cai, Yulong Li, Heather Bradshaw, Ken Mackie, Alex Straiker

**Author notes:** Lead contact: Alex Straiker Indiana University 1101 E 10^th^ St. Bloomington, IN 47405.

## Abstract

Though acetaminophen is a ubiquitous analgesic, its mechanism of action remains unknown. Thus, even though acetaminophen causes ∼500 deaths each year in the US it has not been possible to design safer alternatives. Because endocannabinoids may have a role in acetaminophen action, we examined interactions between the two. We now report that acetaminophen inhibits the activity of diacylglycerol lipase α (DAGLα), but not DAGLβ, decreasing production of the endocannabinoid 2-arachidonoyl glycerol. This gave rise to the counterintuitive hypothesis that decreasing endocannabinoid production by DAGLα *inhibition* may be antinociceptive in certain settings. Supporting this hypothesis, we find that DAGL inhibition by RHC80267 is antinociceptive in wildtype but not CB1 knockout mice in the hotplate test. We propose 1) that activation of DAGLα may exacerbate some forms of nociception and 2) a novel mechanism for the antinociceptive actions of acetaminophen whereby acetaminophen inhibits a DAGLα/CB1-based circuit that plays a permissive role in at least one form of nociception.

## INTRODUCTION

Acetaminophen is the most commonly used medication for pain and fever relief in the US ^1^ and is widely used throughout the world. Though acetaminophen was first synthesized in 1877, its mechanism of action remains surprisingly unclear. Acetaminophen is commonly held to act by selectively inhibiting the enzyme cyclooxygenase 2 (COX2) in the CNS (reviewed in ^2^) but acetaminophen requires high (i.e. millimolar) concentrations to act in this way ^3^. Acetaminophen has also been proposed to act on the cannabinoid signaling system via a secondary metabolite^4,5^.

The endocannabinoid signaling system consists of cannabinoid receptors, lipophilic endocannabinoids (eCB) and the enzymatic machinery to produce and break down these lipid messengers ^6^. Cannabinoid receptors are widely distributed in the CNS and elsewhere in the body. Several studies have linked acetaminophen to activation of the components of the endocannabinoid signaling system. Analgesia induced by acetaminophen requires CB1 receptors ^4,7^. Furthermore, the acetaminophen secondary metabolite N-arachidonoylaminophenol (AM404) has multiple effects on cannabinoid signaling including blockade of anandamide uptake ^8^, inhibition of fatty acid amide hydrolase (FAAH) an enzyme that metabolizes the endocannabinoid arachidonoylethanolamine (AEA, anandamide) ^9^ and indirect activation of TRPV1 ^10^ and/or cannabinoid receptors ^11,12^. However, the concentrations of AM404 achieved in the body after treatment with acetaminophen appear to fall well below of those required to explain these effects (reviewed in ^2^).

While endocannabinoids can produce antinociception, it is often overlooked that knockouts for the 2-AG synthetic enzyme diacylglycerol lipase a (DAGLα) have a strong antinociceptive phenotype in a hot-plate test ^13^. This observation and others^14^ ^15,16^, suggest that inhibiting either endocannabinoid production or CB1 activation may elicit antinociception, and that the antinociceptive properties of endocannabinoids are circuit-specific. We therefore critically examined the interactions of acetaminophen with various steps of endocannabinoid signaling using autaptic hippocampal neurons. In the autaptic model, a single neuron expresses a full retrograde cannabinoid circuit that includes receptor- and depolarization-stimulated postsynaptic synthesis of the endocannabinoid 2-AG by DAGLs ^17^, presynaptic CB1 receptors ^18,19^ and the presynaptic machinery to metabolize 2-AG ^20^. We tested acetaminophen in this model as well as in direct biochemical assays of DAGL activity. We additionally compared the effects of acetaminophen and DAGL inhibition on nociception. Our findings, presented below, suggest an additional model for acetaminophen antinociception by inhibition of DAGLα.

## RESULTS

### Acetaminophen inhibits depolarization induced suppression of excitation in autaptic hippocampal neurons

To test for a direct effect of acetaminophen on excitatory neurotransmission in autaptic hippocampal neurons we performed a series of electrophysiological measurements. The membrane potential of the patched neuron was held at -70 mV and excitatory postsynaptic currents (EPSCs) were elicited every 20 sec with a 1 msec depolarization. Increasing concentrations (10 µM, 30 µM, 100 µM) of acetaminophen did not alter EPSC amplitudes (Figure 1a, relative EPSC charge (1.0 = no change) with acetaminophen (10 μM): 0.98 ± 0.03, n=6, p=0.45; (30 μM): 1.02 ± 0.03, n=7, p=0.34; (100 μM): 1.04 ± 0.04, n=8, p=0.43, NS by one-sample t-test vs. 1.0), indicating that acetaminophen has no direct effect on neurotransmission in our model system.

**Figure 1.**
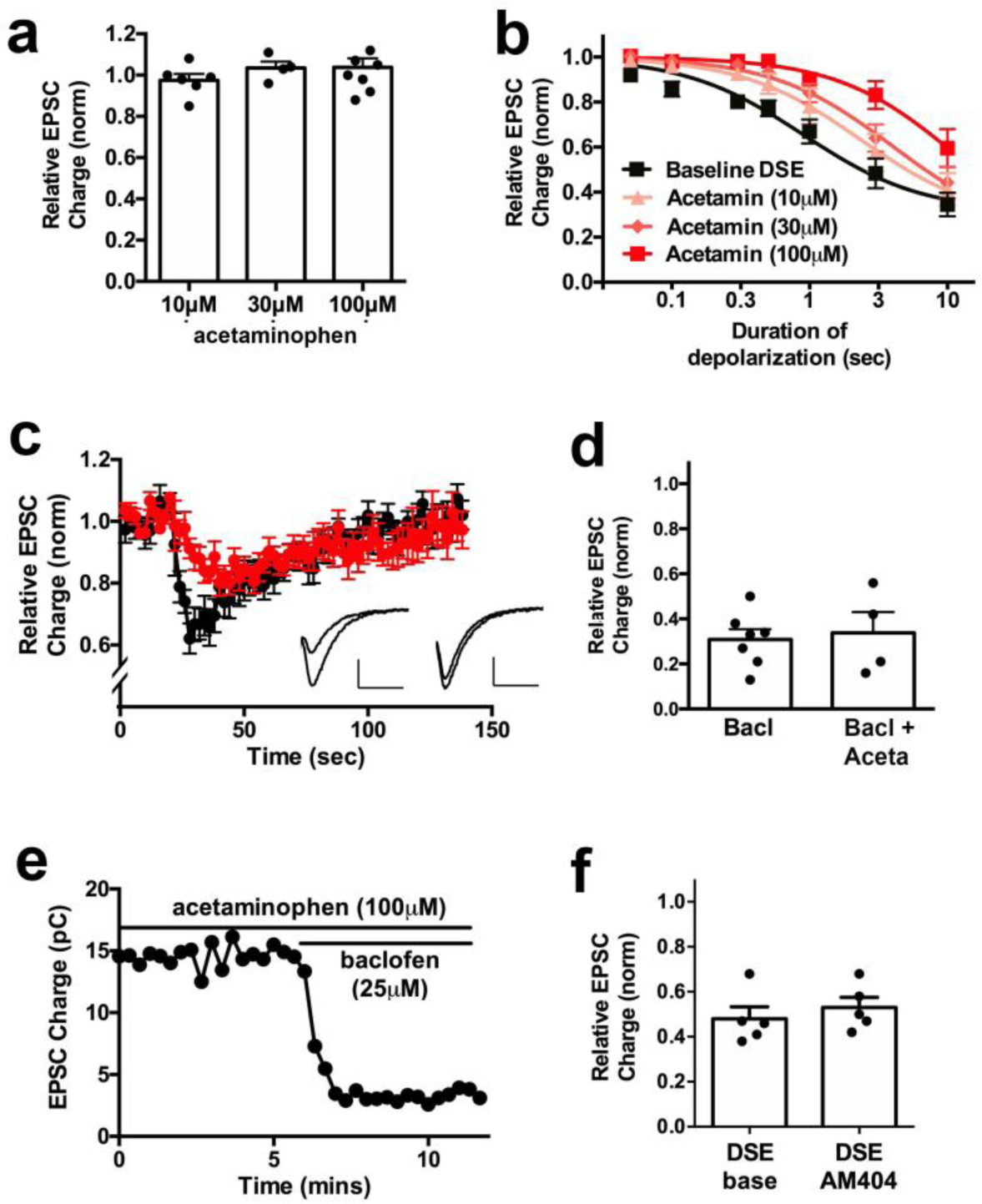
Acetaminophen (APAP) inhibits endogenous cannabinoid neuroplasticity in autaptic hippocampal neurons. A) Acetaminophen does not alter excitatory neurotransmission as measured by evoked EPSC charge. B) Acetaminophen inhibits endocannabinoid-mediated depolarization induced suppression of excitation (DSE) in a concentration-dependent manner. C) Sample time courses of DSE in response to a 3 second depolarization before and during 100 μM acetaminophen. Inset shows representative EPSC traces for control (left) and 100μM acetaminophen (right) after a 3 second depolarization (scale bars: 1 nA and 5 min) D) Responses to the GABA-B agonist baclofen (25 μM) are not altered by 100 μM acetaminophen pretreatment. E) Sample time course for baclofen inhibition of EPSCs in presence of acetaminophen (100 μM). F) AM404 (500 nM) does not affect the magnitude of DSE.

However, we found that acetaminophen inhibited depolarization induced suppression of excitation (DSE), a form of a CB1-mediated retrograde signaling present in autaptic hippocampal neurons. As shown in Figure 1b, acetaminophen inhibits DSE in a concentration-dependent manner, with 10 μM, 30 μM and 100 μM significantly shifting the depolarization-response curves to the right (Figure 1b, ED50 baseline (95% CI): 0.90 sec (0.5-1.6); acetaminophen (10 μM): 2.3 sec (1.9-2.8); acetaminophen (30 μM): 4.0 sec (2.8-5.7); acetaminophen (100 μM): 11.6 sec (4.7-28.5); n=8 for each group, 95% CIs for all concentrations non-overlapping vs. baseline). Averaged DSE responses after 3 sec depolarization before and during 100 μM acetaminophen treatment are shown in Figure 1c. Of note, during acetaminophen treatment DSE both developed more slowly and was reduced in magnitude. Maximal inhibition in response to the longest duration of depolarization was reduced by nearly 40% during treatment with 100μM acetaminophen (EPSC Inhibition after 10 sec depolarization (1.0 = no inhibition, value ± SEM): control: 0.35 ± 0.05; with 100 μM acetaminophen: 0.60 ± 0.08; n=8).

Next, to determine whether acetaminophen was exerting a more general effect by broadly inhibiting G_i/o_ G protein-mediated suppression of neurotransmission, we also tested whether GABA-B signaling was altered by concurrent treatment with acetaminophen (Figure 1d, e). We have previously shown that GABA-B activation robustly inhibits neurotransmission in autaptic neurons in a pertussis-toxin sensitive (i.e., G_i/o_ G protein-mediated) manner ^21^. Here we found that 100 μM acetaminophen did not alter baclofen-induced inhibition of EPSCs (relative EPSC charge with baclofen (25 μM): 0.31 ± 0.05, n=8; baclofen plus acetaminophen (100 μM): 0.34 ± 0.09, n=4; not significant (NS) by unpaired t-test).

We additionally tested whether the acetaminophen metabolite AM404 alters cannabinoid signaling in autaptic neurons. We found that 500 nM AM404 did not alter EPSCs or DSE (Figure 1f: Relative inhibition with DSE (baseline): 0.48 ± 0.04; DSE with AM404 (500 nM, 5 mins): 0.53 ± 0.05; n=5; not significant by paired t-test, p=0.08).

### Acetaminophen does not act presynaptically to inhibit CB1 signaling

The effect of acetaminophen appears to be specific to cannabinoid CB1 receptor signaling. In this case, acetaminophen might be acting presynaptically at CB1 receptors or postsynaptically, by altering synthesis or release of 2-AG. A simple means to distinguish whether acetaminophen is acting pre- or postsynaptically is to test whether acetaminophen alters responses to exogenously applied 2-AG. If acetaminophen acts presynaptically, either at the CB1 receptor itself or some allied protein, then 2-AG responses should be similarly impacted by acetaminophen. If not, then this argues for a post-synaptic effect, likely on eCB synthesis/release. We have previously shown that the EC50 of 2-AG for suppressing EPSCs in autaptic neurons is ∼500 nM and that 5 μM 2-AG produces maximal inhibition. Thus we tested an intermediate 2-AG concentration, 1μM, on its own and in the presence of acetaminophen (100 μM), finding that 2-AG responses were unaltered by acetaminophen (Figure 2; relative EPSC charge with 2-AG (1 μM): 0.54 ± 0.08, n=7; 2-AG plus acetaminophen (100 μM): 0.47 ± 0.08, n=7; NS by paired t-test). This indicates that acetaminophen is likely to act post-synaptically, perhaps by inhibiting synthesis of 2-AG.

**Figure 2.**
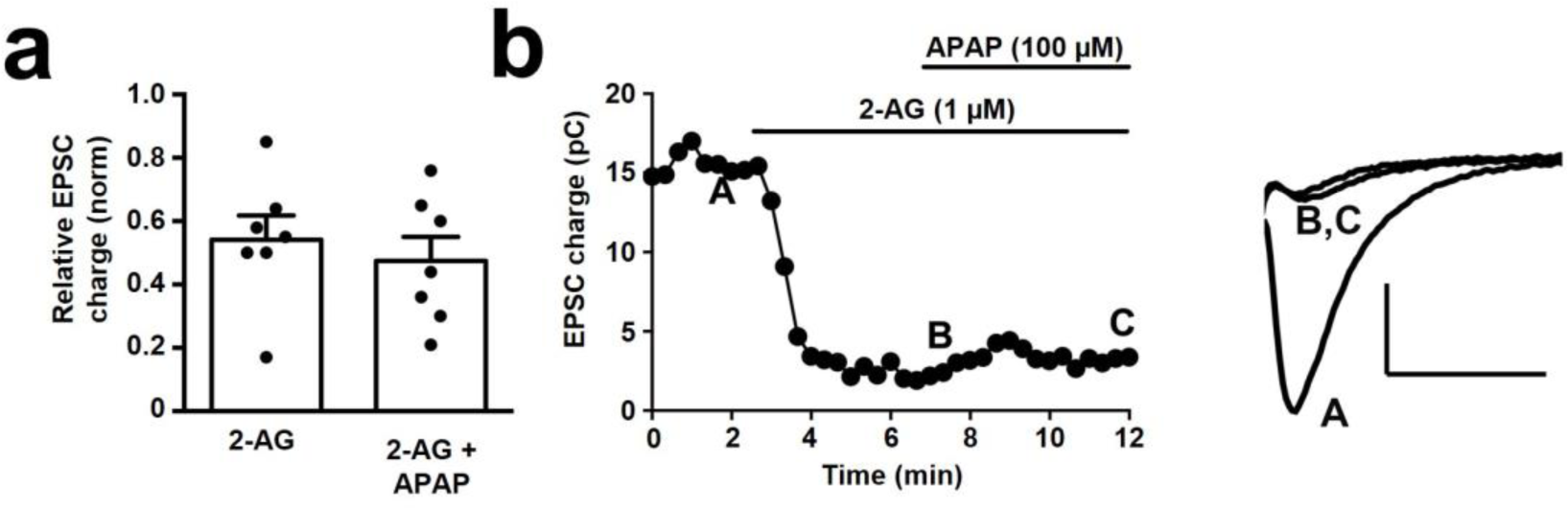
Acetaminophen does not act presynaptically to inhibit CB1 signaling. a) EPSCs are inhibited by ∼40% by 2-AG (1 μM). This inhibition is not reversed by treatment with 100 μM acetaminophen. b) Sample time course shows that 2-AG inhibition of EPSCs is not reversed by acetaminophen treatment. Right panel shows representative EPSC traces for control (a), 2-AG (b) and 2-AG + 100 μM acetaminophen (c) (scale bars: 1 nA and 10 ms).

### Acetaminophen reduces 2-AG synthesis in DAGLα-HEK cells

Diacylglycerol lipases (DAGLs) α and β have been identified as the chief synthesizing enzymes for 2-AG (reviewed in ^22^). We have shown that both DAGLs contribute to DSE in autaptic neurons ^17^. Hypothesizing that acetaminophen might impair DAGL-mediated synthesis of 2-AG, we made use of a human embryonic kidney cell line stably overexpressing DAGLα (HEK-DAGLα). We first treated these cells with 1-stearoyl-2-arachidonoyl-sn-glycerol (S-AG), a compound that has been shown to serve as a substrate for DAGL-mediated 2-AG synthesis ^23^, and compared the resultant lipid profile relative to vehicle-treated cells. Of the examined lipids, 2-AG, 2-LG, and 2-OG, only levels of 2-AG were increased, as expected (Figure 3). Indeed, treatment with 25 μM S-AG resulted in a >40-fold increase in 2-AG levels (results shown in table 1). We also tested for alterations in levels of a panel of acylethanolamines (AEA, PEA, OEA, LEA) but saw no changes after S-AG treatment. We next tested whether acetaminophen (100 μM) altered baseline levels of the above acylglycerols or acylethanolamines, finding that it did not (data not shown). We then tested whether acetaminophen and the DAGLα inhibitor DO34 impacted the S-AG mediated increase in 2-AG and other acyl-glycerols and acylethanolamines, finding that acetaminophen reduced 2-AG levels by about 65% and DO34 reduced 2-AG levels by 70% (Figure 4; nmoles/g of 2-AG after S-AG (25 μM)). Both treatments also decreased levels of 2-OG, and DO34 also decreased 2-LG. All other compounds were not significantly affected by the treatment (results shown in table 2).

**Figure 3.**
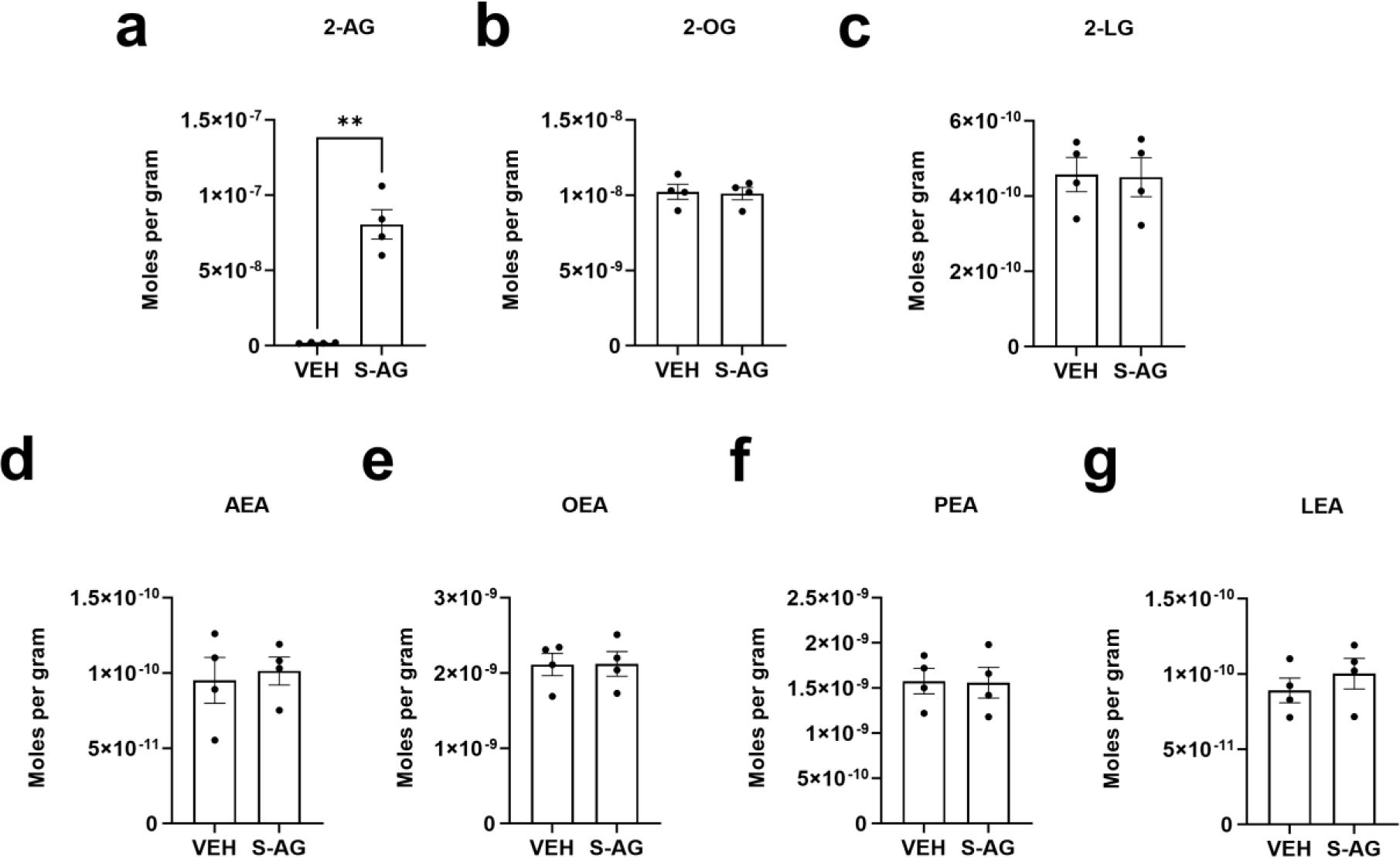
S-AG treatment of HEK-DAGLα cells selectively increases 2-AG levels. Levels of 2-AG (a), 2-OG (b), 2-LG (c), AEA (d), OEA (e), PEA (f) and LEA (g) are shown for HEK-DAGLα cells treated with S-AG (25 μM) or vehicle.

**Figure 4.**
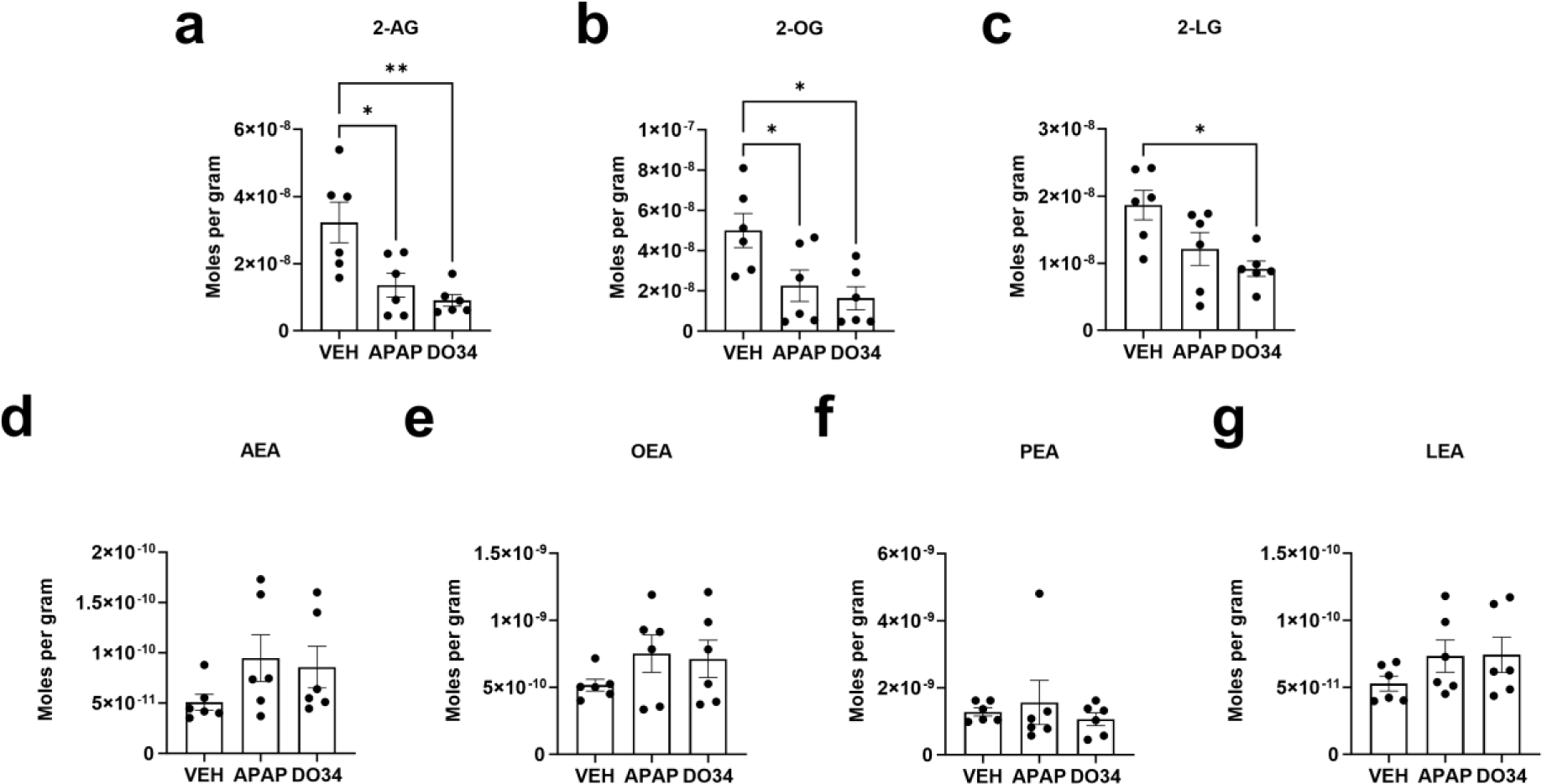
Both acetaminophen and DO34 reduce 2-AG production in HEK-DAGLα cells. Levels of 2-AG (a), 2-OG (b), 2-LG (c), AEA (d), OEA (e), PEA (f) and LEA (g) are shown for HEK-DAGLα cells treated with S-AG (25 µM) + vehicle (VEH), S-AG (25 µM) + Acetaminophen (APAP, 100 µM) or S-AG (25 µM) + DO34 (DO34, 1 µM). *, p<0.05, **, p<0.01 by 1 way ANOVA.

**Table 1.** S-AG increases levels of 2-AG but not other acylglycerols or acylethanolamines examined.

| Investigated lipid | Treatment | nmoles per gram | Standard error | p | N |
| --- | --- | --- | --- | --- | --- |
| 2-AG | VEH | 1.80 | 0.17 | **0.0040** | 4 |
|  | S-AG | 80.56 | 9.78 |  | 4 |
| 2-OG | VEH | 10.22 | 0.50 | 0.8677 | 4 |
|  | S-AG | 10.10 | 0.41 |  | 4 |
| 2-LG | VEH | 0.46 | 0.05 | 0.9196 | 4 |
|  | S-AG | 0.45 | 0.05 |  | 4 |
| AEA | VEH | 0.10 | 0.02 | 0.7431 | 4 |
|  | S-AG | 0.10 | 0.01 |  | 4 |
| OEA | VEH | 2.12 | 0.15 | 0.9740 | 4 |
|  | S-AG | 2.12 | 0.16 |  | 4 |
| PEA | VEH | 1.58 | 0.14 | 0.9481 | 4 |
|  | S-AG | 1.56 | 0.17 |  | 4 |
| LEA | VEH | 0.09 | 0.01 | 0.4292 | 4 |
|  | S-AG | 0.10 | 0.01 |  | 4 |

**Table 2.** Results from the lipid analysis of acylglycerols and acylethanolamines after treatment with S-AG followed by acetaminophen (APAP) and DO34.

| Investigated lipid | Treatment | nmoles per gram | Standard error | p | 1 way ANOVA, F (DFn, DFd) | N |
| --- | --- | --- | --- | --- | --- | --- |
| 2-AG | VEH | 32.27 | 6.04 | **0.0031** | 8.673 (2, 15) | 6 |
|  | APAP | 13.60 | 3.57 |  |  | 6 |
|  | DO34 | 9.03 | 1.76 |  |  | 6 |
| 2-OG | VEH | 50.02 | 8.46 | **0.0137** | 5.794 (2, 15) | 6 |
|  | APAP | 22.56 | 7.83 |  |  | 6 |
|  | DO34 | 16.38 | 5.76 |  |  | 6 |
| 2-LG | VEH | 18.68 | 2.20 | **0.0135** | 5.818 (2, 15) | 6 |
|  | APAP | 12.11 | 2.46 |  |  | 6 |
|  | DO34 | 9.16 | 1.14 |  |  | 6 |
| AEA | VEH | 0.05 | 0.01 | 0.2387 | 1.578 (2, 15) | 6 |
|  | APAP | 0.09 | 0.02 |  |  | 6 |
|  | DO34 | 0.09 | 0.02 |  |  | 6 |
| OEA | VEH | 0.52 | 0.04 | 0.3389 | 1.164 (2, 15) | 6 |
|  | APAP | 0.75 | 0.14 |  |  | 6 |
|  | DO34 | 0.71 | 0.14 |  |  | 6 |
| PEA | VEH | 1.28 | 0.12 | 0.6853 | 0.3876 (2, 15) | 6 |
|  | APAP | 1.56 | 0.66 |  |  | 6 |
|  | DO34 | 1.06 | 0.19 |  |  | 6 |
| LEA | VEH | 0.05 | 0.01 | 0.3089 | 1.272 (2, 15) | 6 |
|  | APAP | 0.07 | 0.01 |  |  | 6 |
|  | DO34 | 0.07 | 0.01 |  |  | 6 |

### Acetaminophen reduces the activity of DAGLα

In a separate assay we tested the effects of acetaminophen on DAGLα or DAGLβ activity in cell lysates using the Enzchek lipase substrate. This assay measures the activity of a broad range of lipases by producing a fluorescent product; activity is measured as accumulated fluorescence over time. We tested the Enzchek substrate on lysates from HEK293 cells transfected with DAGLα (HEK-DAGLα) vs. untransfected control cells (HEK-Ctrl). 30 μM acetaminophen reduced lipase activity in HEK-DAGLα cells, but not in HEK-Ctrl cells or DAGLβ-HEK cells (Figure 5; lipase activity in the presence of acetaminophen in WT HEK cells (1.0 = no effect): 0.89 ± 0.06, n=4; in DAGLα-HEK: 0.76 ± 0.04, n=5; in DAGLβ-HEK: 0.96 ± 0.05, n=4; p<0.05 for DAGLα-HEK by one-sample t-test vs. 1.0). By comparing the average lipase activity of HEK-DAGLα vs. HEK-Ctrl cultures, we calculate that the transfected DAGLα activity amounts to 25% of the total lipase activity in HEK-DAGLα cells (data not shown) and that as a consequence, the effect of acetaminophen 30 µM is to reduce DAGLα activity by 45%. The experiment was repeated with CHO-DAGLα cells and normalized to protein concentrations, with the same findings (data not shown).

**Figure 5.**
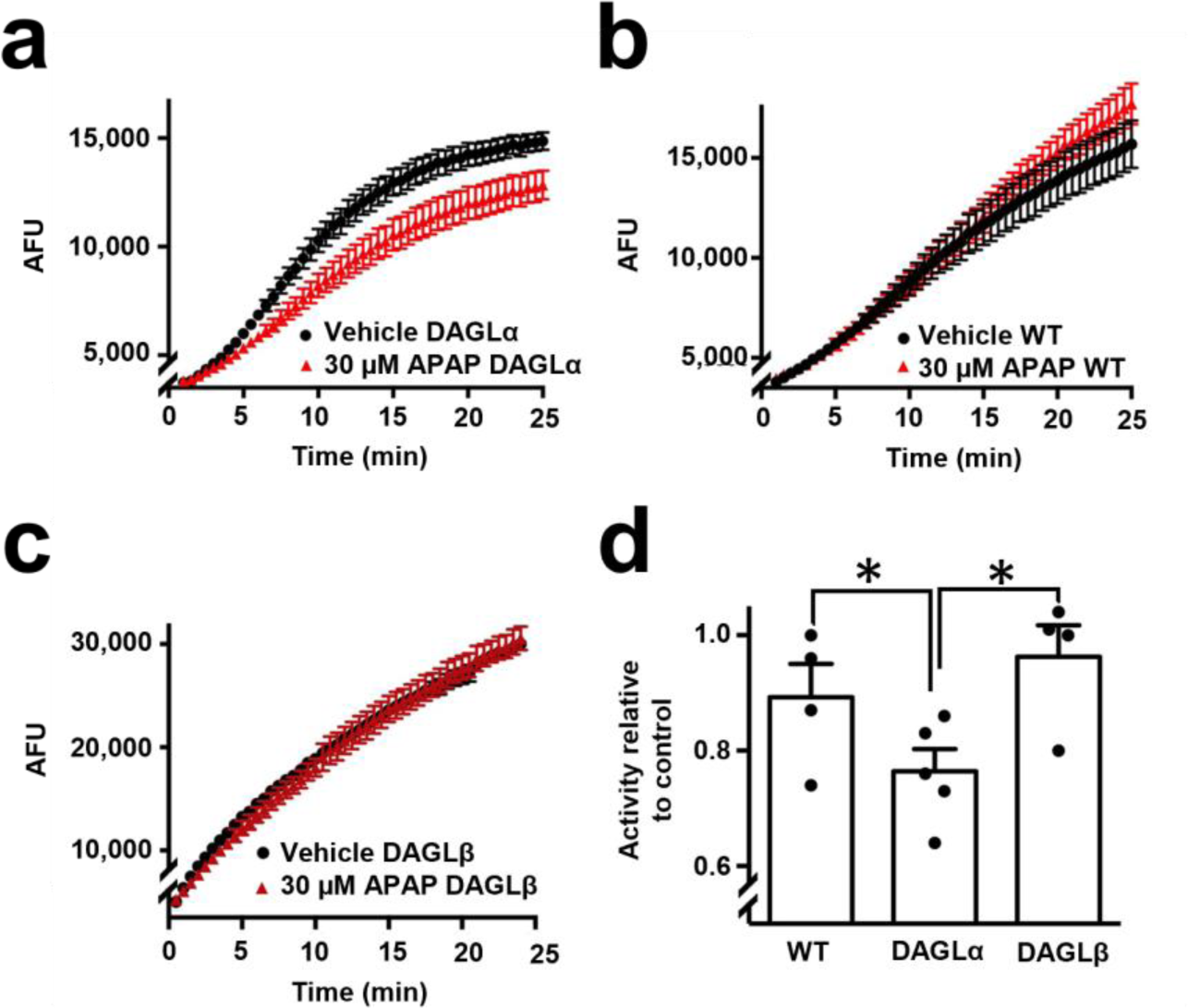
Acetaminophen reduces DAGLα activity. a) A sample time course of Enzchek lipase activity assay in lysates from DAGLα-transfected HEK cells shows that acetaminophen (30 μM) reduces net lipase activity. b) Acetaminophen (30 μM) does not reduce activity in control HEK cell lysates. c) In DAGLβ-transfected cells, acetaminophen does not reduce activity. d) Summary graph showing responses for 30μM acetaminophen in WT, and DAGLα- or DAGLβ-transfected HEK293 cells. *, p<0.05 by unpaired t-test, n=4-5.

To further investigate if acetaminophen inhibited DAGLβ, we examined whether acetaminophen binds to the DAGLβ active site. The DAGL-binding probe HT-01 preferentially and irreversibly binds to the active site of DAGLβ, forming a green fluorescent product that can be detected on a protein gel ^24^. We found that the DAGLβ blocker KT109 (100 nM) competes with the HT-01 probe, but that acetaminophen does not (Figure 6, 100 μM acetaminophen relative to vehicle: 0.94 ± 0.02, n=4; NS, p=0.09 one tailed t-test vs. 1.0 (no effect)).

**Figure 6.**
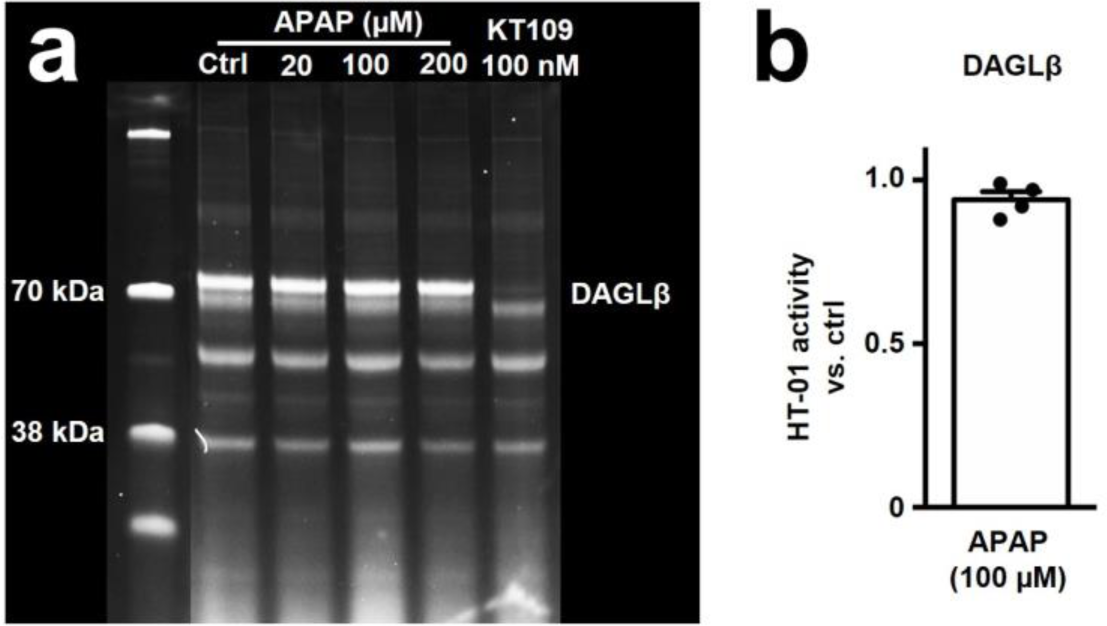
Acetaminophen does not compete with HT-01 for binding to the DAGLβ active site. a) DAGL-binding probe HT-01 is competed by the DAGLβ blocker KT109 (100 nM) but not by increasing concentrations of acetaminophen. b) Summary of results from 4 experiments using 100 µM acetaminophen (APAP). NS, 100 µM treatment, p=0.09 one sample t-test vs. 1.0 (no effect).

### Acetaminophen inhibition of DAGLα-mediated endocannabinoid (eCB) production detected using an eCB sensor

To further examine acetaminophen inhibition of DAGLα we used an endocannabinoid (eCB) sensor. Cells transfected with an eCB sensor readily detect the endocannabinoid 2-AG ^25,26^. In neurons we and others have shown that activation of G_q_-coupled GPCRs elicits production of 2-AG ^19^. We have previously reported that HEK293 cells endogenously express muscarinic M3 receptors ^27^ and hypothesized that activation of these receptors by a muscarinic agonist might induce production of 2-AG. If cells also expressed an eCB sensor then this might offer a real-time read-out of the endogenous synthesis of 2-AG.

HEK293 cells expressing the endocannabinoid sensor ^25,26^ yield a clear fluorescent signal in response to treatment with 2-AG (5 µM) but not to the muscarinic agonist oxotremorine M (oxo-M 10 µM, Figure 7b), consistent with low levels DAGLα. When HEK293 cells are transfected with both eCB3.0 and DAGLα, the cells now respond to both oxo-M and 2-AG, with oxo-M typically producing ∼60% of the maximal 2-AG response (Figure 7a,c).

**Figure 7.**
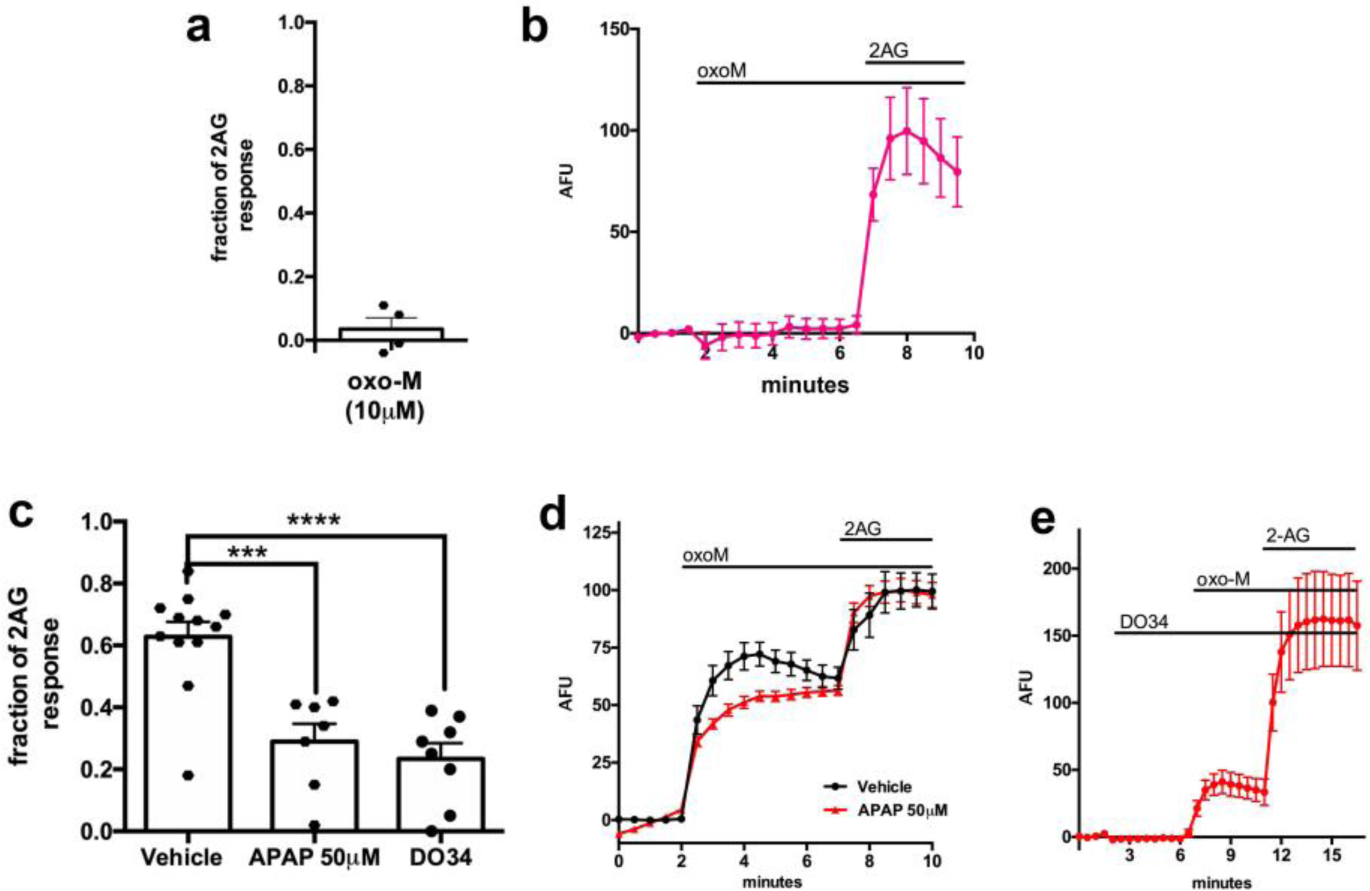
Acetaminophen inhibits muscarinic endocannabinoid production in HEK293 cells co-expressing an eCB sensor and DAGLα. a) Summary of experiments showing muscarinic agonist oxotremorine-M (oxo-M,10 µM) responses relative to 2AG in same cell after vehicle, acetaminophen (APAP, 50 µM) or DO34 (300 nM). b) Sample time course showing that HEK293 cells expressing only eCB3.0 do not respond to 10 µM Oxo-M but robustly respond to 2AG (5 µM). c) summary of experiments showing that acetaminophen (APAP, 50uM) and DO-34 (300nM) each reduce the effect of oxo-M in HEK293 cells co-expressing eCB3.0 and DAGLα. d) Sample time course in HEK293 cells co-expressing eCB3.0 and DAGLα shows responses to oxo-M, 10 µM and subsequent 2-AG. Treatment with acetaminophen (APAP, 50 µM, 10min) reduces this response as a fraction of 2-AG response in the same experiment. Time courses are normalized to 2-AG. e) Sample time course showing effect of DO34 (300 nM) treatment in HEK293 cells co-expressing eCB3.0 and DAGLα. ***, p<0.005, ****, p<0.0001 by 1 way ANOVA with Dunnett’s post hoc test vs. vehicle. n=12, 7, 8

Cells treated with acetaminophen (50 µM, 10 mins) or the DAGLα blocker DO34 (300 nM) exhibit a diminished response to oxo-M (Figure 7a,c-d; vehicle (fraction of max response ± SEM): 0.63 ± 0.04, n=12; acetaminophen(APAP): 0.29 ± 0.06, n=7; DO34: 0.23 ± 0.05, n=8; p<0.005 for APAP vs. Vehicle; p<0.0001 for DO34 vs. vehicle, 1-way ANOVA with Dunnett’s post hoc test vs. vehicle; from at least three independent experiments).

### The antinociceptive effects of acetaminophen: roles for CB1 and DAGL

To learn more about the role of endocannabinoids in the antinociceptive effects of acetaminophen, we employed the hot plate test, a behavioral pain assay that involves supraspinal antinociceptive pathways ^28^ Mallet et al have previously showed that the analgesic effects of acetaminophen in several pain models required the presence of CB1 since they were blocked either by pharmacological inhibition of CB1 or by its absence ^7^. We confirmed this CB1-dependence of acetaminophen analgesia.

Treatment with acetaminophen (300 mg/kg, IP) increased paw withdrawal latencies in wildtype (WT) male and female mice 30 minutes post-injection (Figure 8a-b, differences relative to baseline (± SEM): males, 6.47 ± 0.89 sec, n=6; p=0.0008 two tailed paired t-test; females, 7.28 ± 1.99 sec, n=6; p=0.014 two tailed paired t-test). This antinociceptive effect was lost in male CB**_1_** KO littermates (Figure 8a-b, differences relative to baseline (± SEM): males, -0.19 ± 1.30 sec, n=6; NS, p=0.90 two tailed paired t-test; females: +0.26 ± 1.14 sec, n=6, NS, p=0.82 two tailed paired t-test), consistent with previously reported findings ^7,12^.

**Figure 8.**
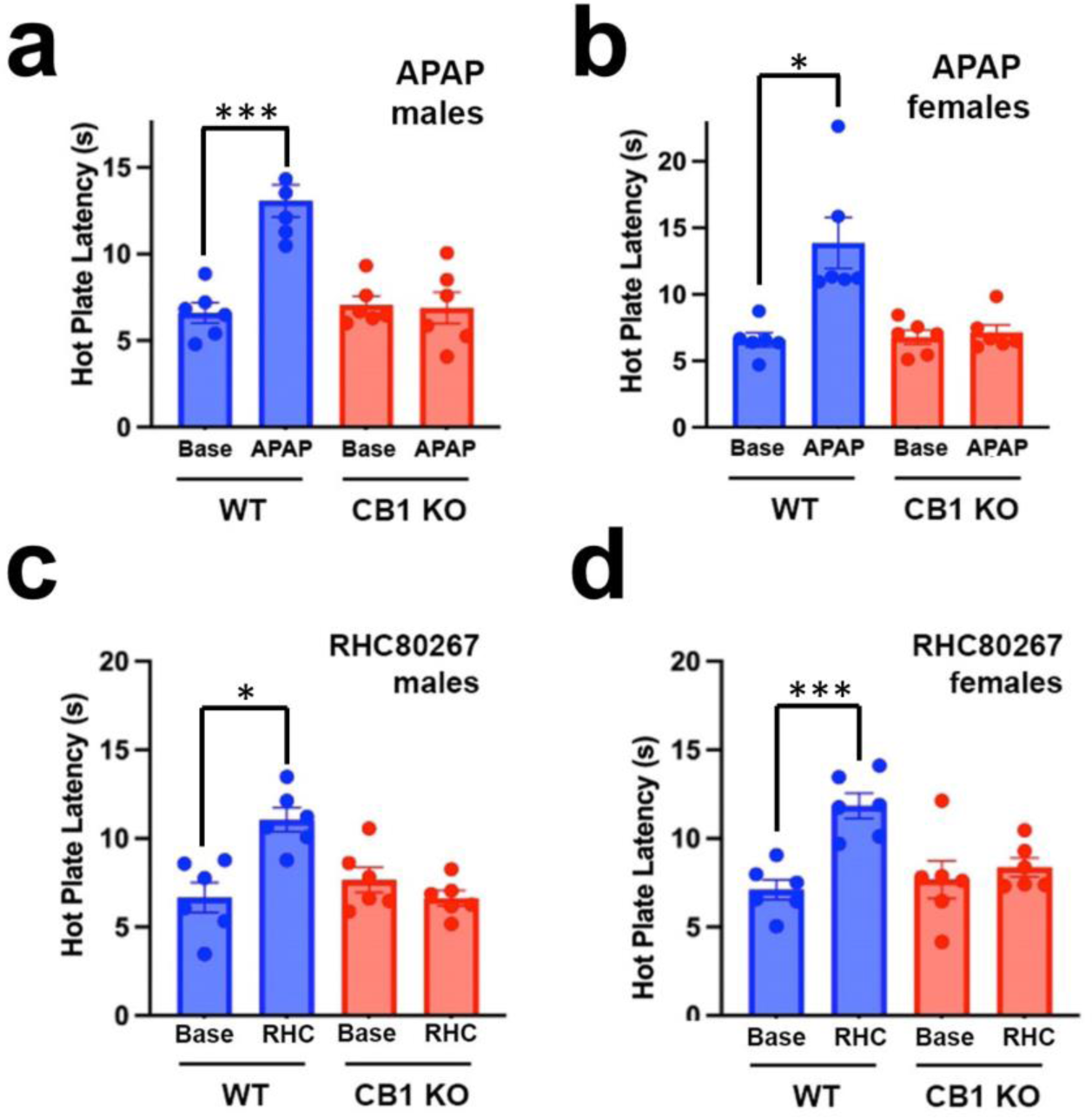
Acetaminophen and the DAGL inhibitor RHC-80267 are antinociceptive in wildtype (WT) but not CB1 knockout mice. a) Acetaminophen (300 mg/kg; i.p.) increased paw withdrawal latency in the hot plate assay relative to baseline in WT but not CB1 KO male mice. b) Same experiment as in a for female mice. c) DAGLα/b inhibitor, RHC-80267 (20 mg/kg i.p.), prolonged the paw withdrawal latency in WT but not CB1 KO male mice. d) Same experiment as in c for female mice. *, p<0.05, ***, p<0.001 by paired t-test vs. pre-drug baseline, n=6 mice per group.

As noted above, DAGLα knockout mice have an antinociceptive phenotype ^13^. Therefore, we tested for potential antinociceptive effects of the DAGLα inhibitor, RHC-80267, in our hot plate assay. We found that RHC-80267 (20 mg/kg, IP) similarly increased paw withdrawal latencies in WT mice, 30 minutes following its administration (Figure 8c-d, differences relative to baseline: males, 4.39 sec ± 1.21, n=6; p=0.015 two tailed paired t-test; females, 4.73 ± 0.59, n=6; p=0.0005 two tailed paired t-test). This increase in paw withdrawal latencies was lost in our CB_1_ KO CD1 mice (Figure 8c-d, differences relative to baseline: males, -1.03 ± 0.48, n=6; NS, p=0.09 two tailed paired t-test; females, 0.69 ± 0.84, n=6; NS, p=0.44 two tailed paired t-test). To test for an effect of acetaminophen on the anandamide synthesis pathway, we additionally tested whether the effects of acetaminophen would be diminished in NAPE-PLD knockout mice, finding that these and WT mice responded similarly: (NAPE-PLD baseline: 7.14 ± 0.67 sec; 300 mg/kg APAP: 11.83 ± 1.03 sec, p=0.0008 by paired t-test vs. baseline, n=9 (5 females, 4 males)).

## DISCUSSION

Acetaminophen is a widely used pain reliever that was first synthesized and characterized in the latter half of the 19^th^ century ^29^. Now, well into the 21^st^ century, acetaminophen’s mechanism of action remains a mystery. Our chief findings are that acetaminophen inhibits cannabinoid signaling in a neuronal model of endogenous cannabinoid neuroplasticity, and that this likely occurs via inhibition of DAGLα-mediated synthesis of the endocannabinoid 2-AG. This effect is seen at physiologically relevant concentrations, well within the therapeutic range of plasma concentrations for acetaminophen (5-20 μg/mL (33-133 μM) ^30,31^). Brain levels of acetaminophen are ∼40% of plasma levels ^32^. We confirmed published findings that the antinociceptive effects of acetaminophen require CB1 receptors ^7^ and we additionally reported that acute inhibition of DAGL is antinociceptive in the hotplate assay and that this also requires CB1 receptors. Based on these findings we propose a model for the antinociceptive effects of acetaminophen. According to this model (depicted schematically in Figure 9), acetaminophen inhibits DAGLα synthesis of the endocannabinoid 2-AG, thereby reducing CB1 activation in an as-yet undefined pro-nociceptive circuit ^14^.

**Figure 9.**
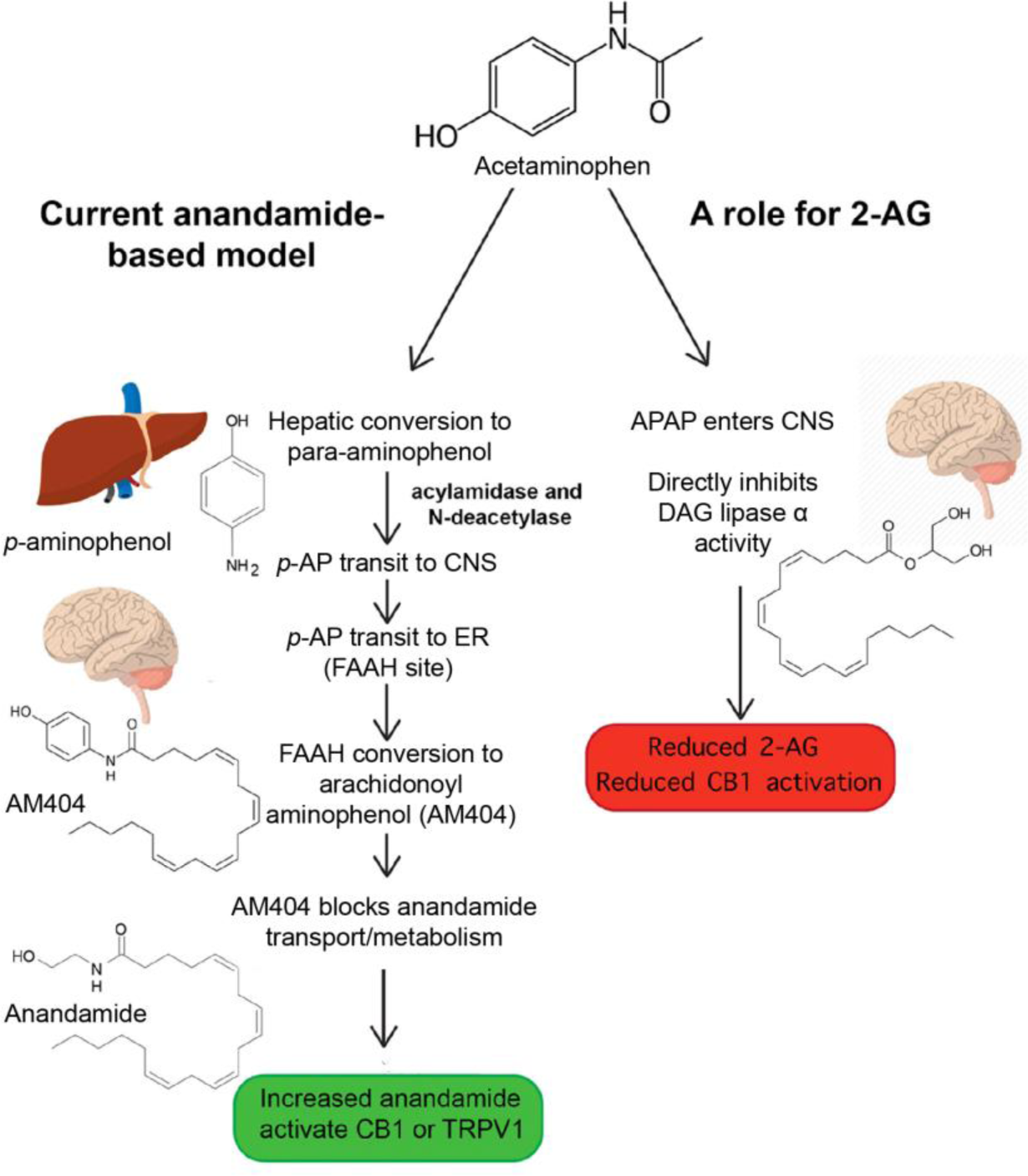
Proposed role for DAGLα and 2-AG in nociception. A schematic depictions of current and proposed acetaminophen (APAP) actions.

Acetaminophen stands out in the complex landscape of pharmacological pain management. On the one hand, it lacks the anti-inflammatory effects – and associated side-effects – of NSAIDs, but its pain-relieving profile is more limited than the powerful opiates. Scientists have long puzzled over how acetaminophen brings about its effects. An early study showed that acetaminophen is a potent inhibitor of prostaglandin E_2_ synthesis, particularly in brain tissue ^33^ and acetaminophen has been proposed to inhibit cyclooxygenase (COX) enzymes ^2^. But acetaminophen is a weak inhibitor of COX1 or COX2, requiring millimolar concentrations ^3^. Acetaminophen has also been proposed to activate the endocannabinoid signaling system via the secondary metabolite arachidonoylaminophenol (AM404) ^9^. We did not observe an effect of AM404 on DSE in the autaptic model of cannabinoid signaling, but the rationale is that acetaminophen is converted to AM404 via a multi-step enzymatic process ^5^ and that AM404 then inhibits anandamide reuptake or metabolism (likely via inhibition of FAAH) ^12,34^, raising anandamide levels and so activating one or more cannabinoid receptors (CB1, CB2 or perhaps TRPV1 ^10^). This rationale is supported by Mallet et al. demonstrating an absence of anti-nociceptive effect of acetaminophen in FAAH and TRPV1 knockout mice ^35^. However, these knockout mice developed with altered endocannabinoid tone, raising the possibility of developmental compensation. According to the mechanism, initially outlined by Hogestatt ^5^, two enzymes (acylamidase and N-deacetylase) sequentially convert APAP to *p*-aminophenol, presumably doing so in the liver. *p*-aminophenol is then hypothesized to travel through the bloodstream, transiting through the blood brain barrier into the CNS, where it is then converted by the enzyme FAAH to AM404. Despite high levels of hepatic FAAH ^34^, significant metabolism of *p*-aminophenol to AM404 appears to occur in the CNS as it is undetectable in blood plasma after acetaminophen treatment ^5^. Because FAAH expression is typically seen intracellularly, likely in endoplasmic reticulum, this calls for an additional transport of p-aminophenol into neurons. This long travel time is potentially problematic since *p*-aminophenol may react with components in bloodstream (e.g. ^36^), or the CNS, raising the question of how much would reach FAAH-expressing cells. And if it is the case that active chaperones in the form of fatty acid binding proteins are needed to escort endocannabinoids to the ER for metabolism by FAAH ^37^ then may require an additional step, as well as time, to similarly transport AM404. The requirement of three enzymes, the circuitous journey for *p*-aminophenol, and that the bulk (>80%) of APAP is metabolized via conjugation of sulfate or glucuronide groups (reviewed in ^38^), may explain why the CNS concentration of AM404 after acetaminophen treatment is insufficient to explain acetaminophen’s effects (reviewed in ^2^). For example, a study of the rate and time course of acetaminophen metabolism to AM404 after a 20 mg/kg dose of acetaminophen reported levels of AM404 in rat brain peaking at 150 pg/g^39^. Such a concentration of AM404 distributed evenly in brain would be <1 nM. In contrast, the concentration of AM404 required for inhibition of FAAH activity in rat brain homogenate was reported to be in the low micromolar range (IC50 3 μM ^9^), as was the reported IC50 for AM404 inhibition of the hypothesized anandamide transporter (IC50, 2 µM ^40^). It is therefore reasonable to ask whether AM404 achieves concentrations in the brain sufficient to explain the hypothesized cannabinoid-dependent effects elicited by inhibition of FAAH or endocannabinoid transport or action on TRPV1 by AM404. Thus, we are left without a clear mechanism of action for acetaminophen. Though the endocannabinoid system is implicated – acetaminophen is without effect in several pain models when CB1 receptors are blocked or deleted ^7^ -- a clue is the observation that knockout mice for DAGLα have a strong antinociceptive phenotype. This combined with the findings presented here has given rise to the model that we have proposed for at least some of the antinociceptive effects of acetaminophen. According to this model, acetaminophen inhibits DAGLα synthesis of the endocannabinoid 2-AG, thereby reducing activity of CB1 receptors (Figure 9). APAP selectively inhibits DAGLα over DAGLβ (Figures 5,6) and APAP analgesia does not require NAPE-PLD.

It is possible that this mechanism was overlooked precisely because many studies find activation of CB1 or CB2 receptors reduces pain or inflammation (reviewed in ^41,42^). However, there is evidence that the reverse can be true: that CB1 activation can be pro-algesic or that CB1 antagonism can be analgesic ^14,15,43,44^ but the evidence in favor of an antinociceptive role for CB1 activation is broad and persuasive. A parsimonious explanation for this duality of the role of eCBs in nociception follows from the ubiquity of the eCB system and the many modulatory pathways that affect nociceptive processing. For example, if CB1 receptors are expressed on a circuit whose activation inhibits a nociceptive pathway (e.g., inhibitory neurons), activation of these receptors will decrease inhibition and increase nociceptive signaling. Conversely, inhibition of these CB1 receptors, either by decreasing 2-AG production or CB1 receptor antagonism will decrease nociceptive signaling. Indeed, such a pathway has been described in the spinal cord ^14^ and likely similar pathways exist elsewhere in the brain. Some of the difference may derive from the puzzling duality of eCB messengers. Anandamide and 2-AG are structurally similar but differ greatly in terms of their metabolism ^45^. The evidence for anandamide and the metabolizing enzyme FAAH nociception are strong, and FAAH KO mice have long been known to have an anti-nociceptive phenotype ^34,46^. Anandamide- and FAAH-dependent antinociception, perhaps in different or parallel circuits affecting pain pathways, may occlude our hypothesized pro-nociceptive DAGLα/CB1 circuit and so explain the observation that acetaminophen is not nociceptive in FAAH KO mice ^47^.

The location and ‘wiring’ of the hypothesized circuit(s) remains to be determined but we may draw some conclusions from what is known of cannabinoid signaling in the CNS. 2-AG-mediated signaling in neurons is typically retrograde, with post-synaptic DAGLα releasing 2-AG onto presynaptic CB1 receptors where these receptors generally inhibit neurotransmission through one or more forms of neuronal plasticity ^6^. The duration of these forms of plasticity span a temporal spectrum from seconds to tens of minutes and perhaps (in the case of long-term depression) longer. Lesion studies that implicated cannabinoid signaling in the effects of acetaminophen found roles for components of the descending pain pathways ^7^ so it is possible that we are dealing with a retrograde inhibitory circuit along this pathway.

This model raises some interesting questions about the interaction of acetaminophen with DAGLα. Is it a feature of the particular synapse? Is DAGLα producing 2-AG tonically, or is DAGLα activity induced and if so by what stimulus? Is the DAGLα activity upregulated by painful stimuli? DAGLα activity can be stimulated variously by depolarization and activation of post-synaptic G_q_-coupled metabotropic receptors ^6^. There is also evidence that basal activity of the enzyme can be modulated by intracellular messengers ^23^.

Though the focus of our discussion has been on neuronal actions of CB1, it is important to note the potential contribution of glial or microglial cannabinoid signaling. A growing body of literature points to a glial cannabinoid role as reviewed in ^48^ and DAGLα may be present in astrocytes ^49^. While we have offered evidence for a neuron-based DAGL-dependent role in cannabinoid signaling in autaptic neurons ^17^, those findings do not rule out an astrocyte role in cannabinoid signaling in pathways involved in nociception. Similarly, our results do not rule out the contribution of secondary metabolites such as prostaglandins that may exert effects on nociceptive pathways.

In summary, we have determined that acetaminophen reduces 2-AG production in a neuronal model, likely via inhibition of the eCB-synthesizing enzyme DAGLα. The effect occurs at concentrations that are within the therapeutic range of plasma concentrations of acetaminophen. Moreover, acute inhibition of DAGL is anti-nociceptive in the hotplate test in a CB1-dependent manner. We propose a new model to explain at least some of the the anti-nociceptive effects of acetaminophen, one that involves inhibition of a DAGLα/CB1 neuronal circuit that modulating nociception. This mechanism may be part of a multimodal effect of acetaminophen that includes both through direct action and via the action of metabolites as proposed elsewhere. But our findings point to DAGLα inhibition as a potential novel target for development of therapeutics for pain management.

## ACKNOWLEDGMENTS

This work was funded by R01AT011162 (KM), T32DA024628 (TB), F31DA057835 (TB) and Fulbright fellowship (MD).

## AUTHOR CONTRIBUTIONS

Conceptualization: AS, KM, HB; Methodology: AS, HB; Investigation: MD, TBB, JB, EL, AZ, AG, JWM, AS, NM; Resources: RC, YL, SC; Data analysis: MD, TBB, EL, AS; Supervision: AS, KM, HB; Writing—original draft: AS, MD, KM; Writing—review & editing: AS, MD, KM

## METHODS

## RESOURCE AVAILABILITY

### Lead Contact

Further information and requests for resources and reagents should be directed to and will be fulfilled by the lead contact, Alex Straiker.

### Materials Availability

This study did not generate new unique reagents.

### Data and Code Availability

Any additional information required to reanalyze the data reported in this study is available from the lead contact upon request.

### Data

All primary data reported in this study will be shared by the lead contact upon request.

### Code

This publication does not report original code.

Any additional information required to reanalyze the data reported in this publication is available from the lead contact upon request.

## EXPERIMENTAL MODEL AND SUBJECT DETAILS

### Animals

All procedures used in this study were approved by the Animal Care Committee of Indiana University and conform to the Guidelines of the National Institutes of Health on the Care and Use of Animals. Mice were bred and group-housed in accordance with animal welfare rules in a pathogen-free facility with temperature 22 ± 2°C, 45% humidity, 12:12-h light/dark cycle, and food and water *ad libitum*. For behavioral testing wild type and CB1 knockout mice on a CD1 background strain of either sex, aged 3-5 months were used. NAPE-PLD knockout mice were on a C57BL/6 background.

### Hippocampal neurons culture

Mouse hippocampal neurons isolated from the CA1-CA3 region were cultured on microislands as described previously ^50,51^. Neurons were obtained from animals (age postnatal day 0-2, undetermined sex) and plated onto a feeder layer of hippocampal astrocytes that had been laid down previously ^52^. Cultures were grown in high-glucose (20 mM) DMEM containing 10% horse serum, without mitotic inhibitors and used for recordings after 8 days in culture and for no more than three hours after removal from culture medium.

## METHOD DETAILS

### Electrophysiology

When a single neuron is grown on a small island of permissive substrate, it forms synapses or “autapses” onto itself. All experiments were performed on isolated autaptic neurons. Whole cell voltage-clamp recordings from autaptic neurons were carried out at room temperature using an Axopatch 200A amplifier (Molecular Devices, Sunnyvale, CA). The extracellular solution contained (in mM) 119 NaCl, 5 KCl, 2.5 CaCl_2_, 1.5 MgCl_2_, 30 glucose, and 20 HEPES. Continuous flow of solution through the bath chamber (∼2 ml/min) ensured rapid drug application and clearance. Drugs were typically prepared as stocks, and then diluted into extracellular solution at their final concentration and used on the same day.

Recording pipettes of 1.8-3 MΩ were filled with (in mM) 121.5 KGluconate, 17.5 KCl, 9 NaCl, 1 MgCl_2_, 10 HEPES, 0.2 EGTA, 2 MgATP, and 0.5 LiGTP. Access resistance and holding current were monitored and only cells with both stable access resistance and holding current were included for data analysis. Conventional stimulus protocol: the membrane potential was held at –70 mV and excitatory postsynaptic currents (EPSCs) were evoked every 20 seconds by triggering an unclamped action current with a 1.0 ms depolarizing step. The resultant evoked waveform consisted of a brief stimulus artifact and a large downward spike representing inward sodium currents, followed by the slower EPSC. The size of the recorded EPSCs was calculated by integrating the evoked current to yield a charge value (in pC). Calculating the charge value in this manner yields an indirect measure of the amount of neurotransmitter released while minimizing the effects of cable distortion on currents generated far from the site of the recording electrode (the soma). Data were acquired at a sampling rate of 5 kHz.

DSE stimuli: After establishing a 10-20 second 0.5 Hz baseline, DSE was evoked by depolarizing to 0 mV for 50 ms, 100 ms, 300 ms, 500 ms, 1 s, 3 s and 10 s, followed in each case by resumption of a 0.5 Hz stimulus protocol for 20-80+ seconds, allowing EPSCs to recover to baseline values. This approach allowed us to determine the sensitivity of the synapses to DSE induction. To allow comparison, baseline values (prior to the DSE stimulus) are normalized to one. DSE inhibition values are presented as fractions of 1, i.e. a 50% inhibition from the baseline response is 0.50 ± standard error of the mean. The x-axis of DSE depolarization-response curves are log-scale seconds of the duration of the depolarization used to elicit DSE. Depolarization response curves are obtained to determine pharmacological properties of endogenous 2-AG signaling by depolarizing neurons for progressively longer durations (50 ms, 100 ms, 300 ms, 500 ms, 1 s, 3 s and 10 s). The data were fitted with a nonlinear regression (Sigmoidal dose response; GraphPad Prism 6, La Jolla, CA), allowing calculation of an ED50, the effective dose or duration of depolarization at which a 50% inhibition is achieved. A statistically significant differences between these curves is defined as non-overlapping 95% confidence intervals for the ED50s. Values on graphs are presented as mean ± S.E.M.

### HEK293-DAGLα cell line

To derive a stable HEK293-DAGLα cell line, HEK293 cells were transfected with diacylglycerol lipase α (DAGLα) in a pEF6 vector using Lipofectamine 2000 ^17^. Colonies were selected using blasticidin (10 µg/mL). The DAGLα has a V5 epitope inserted on the carboxyl end of the sequence that was used to facilitate the identification cell clones stably expressing DAGLα. Cells were used at passages ∼10-12.

### Lipid analysis methods

Levels of 2-acylglycerols and N-acylethanolamines were measured by HPLC/MS/MS from DAGLα−HEK293 cultures as previously described ^53^. Briefly, HEK293 cells overexpressing DAGLα were grown in separate 10cm petri dishes (i.e., one dish represented one biological replicate) and treated with either vehicle, 25 µM 1stearoyl-2-arachidonoyl-*sn*-glycerol (S-AG), or S-AG plus 100 µM Acetaminophen in serum-free media for 2 hours at 37C. The media solution was aspirated and then cells were treated with 2mls of HPLC-grade methanol, scraped, and transferred to a centrifuge tube, deuterium-labeled standard added, and the cell solution was held on ice and covered for one hour. The cell solution was then sonicated on ice for 1 minute, then centrifuged at 19,000 x g at 24°C for 20 min, and the supernatant was collected. The supernatant was diluted to 20% with HPLC-grade water and then the matrix was partially purified on C18 solid-phase extraction columns (Agilent Technologies, Santa Clara, CA) as previously described ^53^. For data presented in Table 1 and Figure 3, HPLC/MS/MS analysis was performed as previously described using a C18 Zorbax reversed-phase analytical column (Agilent Technologies, Santa Clara, CA) and an API 3000 triple quadrupole mass spectrometer (Applied Biosystems/DSM Sciex, Foster City, CA). Analysis for each lipid was as previously described ^53^. For data presented in Table 2 and Figure 4, samples were analyzed using an API 7500 coupled to a Shimadzu LC system LC-40DX3 (Kyoto, Japan) with a reversed-phase Luna C18(2) HPLC column (Phenomenex, Torrence, CA) as described previously ^54^.

### Enzchek lipid activity assay

Multiple 10 cm plates of HEK cells were grown in parallel. When cells reached ∼70% confluency, half of the plates were transfected with DAGLα or DAGLβ and mcherry. The following day these were compared with untransfected wild-type cells plated on the same day. Membrane proteins were extracted from cells by treating them with 1 ml of a freshly prepared lysis buffer consisting of 50 mM HEPES, 100 mM NaCl, 5 mM CaCl2, 0.5% TritonX-100, pH 7.2. Cells were treated with lysis buffer for 5 minutes and then centrifuged for 5 minutes at 4°C, after which the supernatant was removed and maintained on ice for analysis on the same day. Lipase activity was evaluated on samples in a 96 well plate using a Flexstation III. Wells received protein extract (50 μL); lysis buffer (50μL); drug or vehicle (5 μL). Enzchek (20 μM in 5 μL) was added just before readings were begun (485 nm stimulation/515 nm emission, six readings per time point, 30 s intervals). Samples were run in pairs (technical replicates) in a given experiment. Experiments were repeated from separate cultures (biological replicates). Activity was calculated as the slope of the curve during the linear phase. Experiments were repeated as above using CHO cells transfected with DAGLα.

### Gel-based competitive ABPP experiments

A mouse was sacrificed, its brain removed, rinsed with lysis buffer (50 mM HEPES, 100 mM NaCl, 5 mM CaCl_2_, 0.5% TritonX-100, pH 7.2) and transferred to a Dounce homogenizer with 2 mL of cold lysis buffer and homogenized. The homogenate was maintained on ice for 15 minutes, then transferred to Eppendorf tubes and spun down (1,000 x g, 5 mins, 4°C). The supernatant was taken and protein concentration determined. Samples were analyzed the same day as prepared. For gel-based competitive ABPP experiments, supernatants (1 mg/mL) were treated with drug or vehicle for 30 mins, then treated with HT-01 ^24^, an activity-based probe for DAGLβ activity (1 μM final concentration) in a 50 μL total reaction volume. After 30 min at 37 °C, the reactions were quenched with SDS-PAGE loading buffer (In 10 mL: 2 mL 1M Tris-HCl, pH 6.8; 0.8 g SDS; 4 mL 100% glycerol; 1 mL 0.5M EDTA; 0.2% bromophenol blue; 4% β-mercaptoethanol). After separation by SDS-PAGE (10% acrylamide), samples were visualized by in-gel fluorescence scanning using a Typhoon 9500 fluorescent scanner. Intensity of fluorescent bands was measured using Fiji. To confirm the identity of the HT-01 labeled DAGLβ band, we tested whether signal of the predicted 74 kDa target band was blocked by the DAGLβ blocker KT109 ^24^ (100 nM). Experiments were run from three separate animals and compared to control values using a one-sample t-test.

### A cannabinoid sensor-based assay of endocannabinoid synthesis

HEK293 cells were co-transfected with DAGLα and the eCB sensor ^25,26^ using Lipofectamine 2000 according to the manufacturer’s instructions. Here we used the eCB3.0 sensor, which offers improved properties over the eCB2.0 sensor ^26^. After transfection was visually confirmed, the cells were transferred to 96 well plates and tested the following day. Transfected cells were visualized in a given well and recorded using an inverted Nikon Instruments microscope fitted with a Spectra-X light engine (Lumencor, Beaverton, OR), a Flash 4 Hamamatsu camera (Hamamatsu City, Japan) and Nikon Elements software. At the end of each experiment, cells were treated with 5 µM 2-AG to determine a maximal response. Oxotremorine-M responses were given as a percentage of the maximal 2-AG response. Multiple cells (4-11) were averaged for a single experiment (technical replicates). Cells were prepared from at least three separate batches on separate days for experiments (experimental replicates). Before a given set of experiments, cell media was replaced with ACSF (see Electrophysiology). Drugs were added directly to a given well at 2x, 3x, 4x concentrations as appropriate. Oxo-M responses were calculated at 2 minutes post-treatment.

### Hot plate behavioral experiments

The antinociceptive effects of acute acetaminophen (300 mg/kg) and RHC-80267 (20 mg/kg) were assessed in wildtype and CB_1_ knockout male and female CD1 mice using the hot plate pain test. All mice weighed ∼25-40 g and were ∼14-20 weeks old. Mice were placed on a 56℃ hot plate (IITC Life Science Incremental Hot/Cold Plate) until nociceptive behaviors such as jumping, paw licking, or paw shaking were observed, or until the cut-off latency of 30 seconds was observed. Baseline (pre-injection) responses were measured at least 7 hours prior to drug administration, and post-injection responses were assessed 30 minutes following drug administration. Acetaminophen and RHC-80267 were administered via intraperitoneal (i.p.) injection to mice in a final volume of 10 mL/kg and each drug was dissolved in vehicle consisting of 100% ethanol, Kremophor, and PBS in a 1:1:18 ratio and used the same day for experimentation.

Baseline and post-drug hot plate data were expressed as the mean (± S.E.M) latency in expressing pain behaviors. Post-drug latencies were compared to baseline latencies using a paired t-test. These experiments used an n=6 mice per group.

### Drugs

Acetaminophen was obtained from Tocris-Biosciences (Bristol, UK). S-AG, RHC-80267 and 2-AG were purchased from Cayman Chemical (Ann Arbor, MI). Baclofen was purchased from Sigma-Aldrich (St. Louis, MO).

### Quantification and statistical analysis

GraphPad Prism 6 (La Jolla, CA) software was used for the statistical analysis. Data are presented as means ± SEM. The statistical details of experiments can be found in the results section. Differences in EPSC amplitudes were tested by one-sample t-test. Relative EPSC changes with baclofen were compared by unpaired t-test. The effect of AM404 on DSE and effect of 2-AG on EPSC were tested with paired t-test. The DSE dose-response data were fitted with a curve with variable slope (four parameters) and the ED50 and 95% CI are provided in the results. For the comparison of lipid levels in cells 1 way ANOVA was used. The effect of acetaminophen on DAGLα activity was determined by one-sample t-test. The mouse nociception data were analyzed by two tailed pair t-test. Differences were considered significant at p ≤ 0.05.

### Key resources table

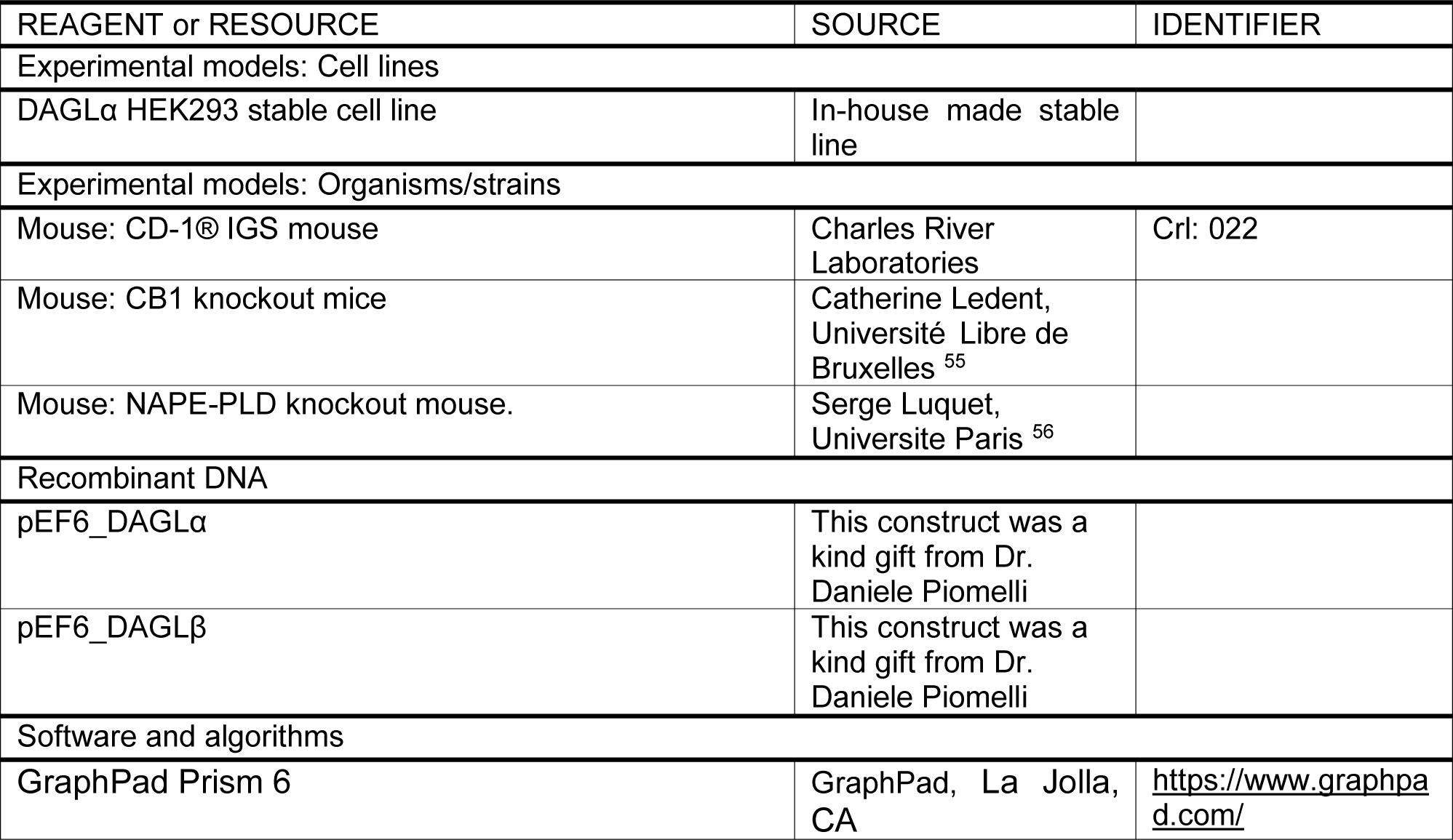

| REAGENT or RESOURCE | SOURCE | IDENTIFIER |
| --- | --- | --- |
| Experimental models: Cell lines |  |  |
| DAGLα HEK293 stable cell line | In-house made stable line | |
| Experimental models: Organisms/strains |  |  |
| Mouse: CD-1® IGS mouse | Charles River Laboratories | Crl: 022 |
| Mouse: CB1 knockout mice | Catherine Ledent, Université Libre de Bruxelles <sup>55</sup> |  |
| Mouse: NAPE-PLD knockout mouse. | Serge Luquet, Université Paris <sup>56</sup> |  |
| Recombinant DNA |  |  |
| pEF6_DAGLα | This construct was a kind gift from Dr. Daniele Piomelli | |
| pEF6_DAGLβ | This construct was a kind gift from Dr. Daniele Piomelli | |
| Software and algorithms |  |  |
| GraphPad Prism 6 | GraphPad, La Jolla, CA | https://www.graphpad.com/ |

